# Three-dimensional tracking of tethered particles for probing nanometer-scale single-molecule dynamics using plasmonic microscope

**DOI:** 10.1101/2021.05.29.446317

**Authors:** Guangzhong Ma, Zijian Wan, Yunze Yang, Wenwen Jing, Shaopeng Wang

**Affiliations:** Biodesign Center for Biosensors and Bioelectronics, Arizona State University, Tempe, Arizona 85287, USA

**Author notes:** Correspondence to: **Correspondence and requests for materials** should be addressed to S.W.

## Abstract

Three-dimensional (3D) tracking of surface-tethered single-particle reveals the dynamics of the molecular tether. However, most 3D tracking techniques lack precision, especially in axial direction, for measuring the dynamics of biomolecules with spatial scale of several nanometers. Here we present a plasmonic imaging technique that can track the motion of ∼100 tethered particles in 3D simultaneously with sub-nanometer axial precision at millisecond time resolution. By tracking the 3D coordinates of tethered particle with high spatial resolution, we are able to determine the dynamics of single short DNA and study its interaction with enzyme. We further show that the particle motion pattern can be used to identify specific and non-specific interactions in immunoassays. We anticipate that our 3D tracking technique can contribute to the understanding of molecular dynamics and interactions at the single-molecule level.

## Introduction

Molecules in biological systems perform their function by traveling between different locations and interacting with other molecules. Tracking the motion of single molecule is of fundamental importance to understand molecular heterogeneity^1^, interactions^1,2^, and myriad intracellular processes^3^. Biomolecules such as protein and DNA are only a few nanometers in size, which requires tracking techniques to have resolution at least comparable to the size to precisely reveal the motion and intramolecular dynamics. On the other hand, the time scale of biomolecular dynamics ranges from microseconds to hours, and thus high temporal resolution and long tracking duration are entailed. Over the past decades, single-molecule fluorescence has emerged as the mainstream technique to track single molecules by incorporating fluorescent dyes into the molecules. However, due to photobleaching and the limited number of photons emitted from a single molecule, the temporal resolution, tracking precision and duration are compromised. To overcome these limitations, nanoparticles are used as an alternative label owing to the strong optical signals. Sub-nanometer precision with microseconds time resolution has been achieved.^4, 5^ However, most fluorescence-and particle-based tracking techniques only track the projection of the motion in the imaging plane (x and y directions) for ease of operation, which may lead to biased results due to missing information in the third dimension (axial or z direction).^6,7^ Measuring the third dimension introduces complexity to the existing 2D tracking system^6, 8^, including additional optical components and data processing complexity. One method to determine particle axial movement is to analyze the size and shape of the off-focus patterns arising from diffraction.^9-11^ Other methods utilize high-speed laser scanning in multiple focal planes to localize the particle and move the objective to follow the particle motion via a feedback system.^12, 13^ However, it is still challenging to achieve sub-nanometer resolution in z-direction at kHz frame rate. There is a need to develop a simple yet precise 3D tracking technique to measure single-molecule dynamics.

Surface plasmon resonance microscopy (SPRM)^14^ is capable of tracking nanoparticles in 3D. Unlike the aforementioned 3D techniques, SPRM extracts axial information directly from the scattering intensity of particles within the evanescent field, which does not introduce additional complexity to the existing system. Since the evanescent field decays exponentially from the surface, the scattering intensity is highly sensitive to z displacement, rendering sub-nanometer resolution in z direction. Together with its nanometer resolution in xy directions and millisecond time resolution, SPRM meets all the requirements for 3D single-molecule dynamics study. Although preliminary studies have demonstrated using SPRM to track single organelle transportation,^15^ mechanical oscillation of nanoparticles,^16^ and thermal fluctuations of nanoparticles tethered by proteins,^2^ its advantage in probing single-molecule dynamics and molecular interactions in 3D still remains to be exploited.

In this work, we present SPRM as a multiplexed 3D single-particle tracking technique with sub-nanometer axial resolution at up to kHz frame rate, which allows further analysis of the dynamics of single molecules attached to the particles. We first use short DNA tethered particles as a model system to demonstrate the capability of 3D tracking and the benefits in comparison with traditional 2D tracking. Then we study the interaction between DNA and a DNA helicase and derive the unwinding rate and rotation angle of the helicase from tracking results. We also show our technique is useful in particle-based immunodetection in terms of identification and removal of non-specific interactions from specific ones. The camera-based detection owns the capability of tracking over 100 individual particles simultaneously, which provides enough throughput to generate statistics in a single measurement. We envisage the high-precision 3D single particle tracking will provide new insight in single molecule detection and biosensing.

## Results

### Detection principle

Particles are tethered to a gold surface using DNA or protein molecules. An objective-based plasmonic imaging setup^17^ is used for tracking the tethered particles (Figure 1a). The surface plasmonic wave is excited on the gold surface using a superluminescent diode (SLED), which is then scattered by particle and generates a bright spot with parabolic tails due to interference (Figure 1b). This pattern is known as the point spread function (PSF) of particle under SPRM. Figure 1b shows the SPR image of over 100 DNA tethered particles that can be tracked simultaneously. The lateral (xy) intensity profile of each single particle shows a Gaussian distribution in x direction and a skewed Gaussian distribution in y direction, respectively (Figures 1c-d and Supplementary Note 1). Therefore, the peak position can be used to localize the particle in xy plane. We localize the particles using a single-particle tracking software, TrackMate,^18^ which automatically finds the local maximum for each particle and tracks the motion over time. After obtaining the local maximum, the mean intensity of all pixels surrounding the local maximum within a circle of ∼1.5 µm diameter is used to calculate the axial position *z*, given by 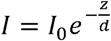, where *I* is the mean image intensity, *I*_0_ is the intensity when the particle is closely attached to the surface (at *z* = 0) and *d* is the decay constant of evanescent field (∼100 nm). To evaluate tracking precision, we recorded the relative movement between two particles stuck on the surface at 100 frames per second (fps) for 6 s and calculated the standard deviation in x, y and z directions. The standard deviation is defined as localization precision, which is ∼2 nm in xy, and 0.44 nm in z (Figure 1e). Due to the exponential decay of evanescent field, localization in z is more precise than xy, which is a unique feature of SPR. We also note that by increasing the incident light intensity, higher localization precision (1 nm in xy and 0.1 nm in z) and temporal resolution (1000 fps) can be achieved (see Discussion and Figure S1). Using the experimental condition and spaciotemporal resolution in Figure 1e, we tracked the motion of a free 1 µm polystyrene (PS) particle near the surface in 3D as an example (Figure 1f). The high resolution revealed detailed nanometer-scale information, including particle-surface interaction (the c-shape pattern) and Brownian motion (the scattered pattern), as shown in Figure 1f. To demonstrate the tracking accuracy, we introduced a second imaging channel additional to the existing SPR channel, which used transmitted light to simultaneously track the particle motion in 2D (see Figure S2 for the setup). The xy projection of the 3D SPR pattern in Figure 1f was similar to the pattern obtained by transmitted light tracking (Figure 1g and Supplementary Movie 1), except for minor deviation. The deviation in x and y directions were quantified respectively by constructing correlation curves using the x and y coordinates determined from the two tracking methods (Figure 1h). The correlation in both directions were strong with *R*^2^ > 0.997, indicating SPR has excellent tracking accuracy. However, the slope of the correlation curves in x and y were 0.908 and 1.17, or -9.2% and 17% deviation, respectively. This deviation is linear because the *R*^2^ is close to 1. Further analysis suggests that the deviation is likely due to fitting algorithm and different imaging principle between SPR and transmitted imaging (see Discussion).

**Figure 1.**
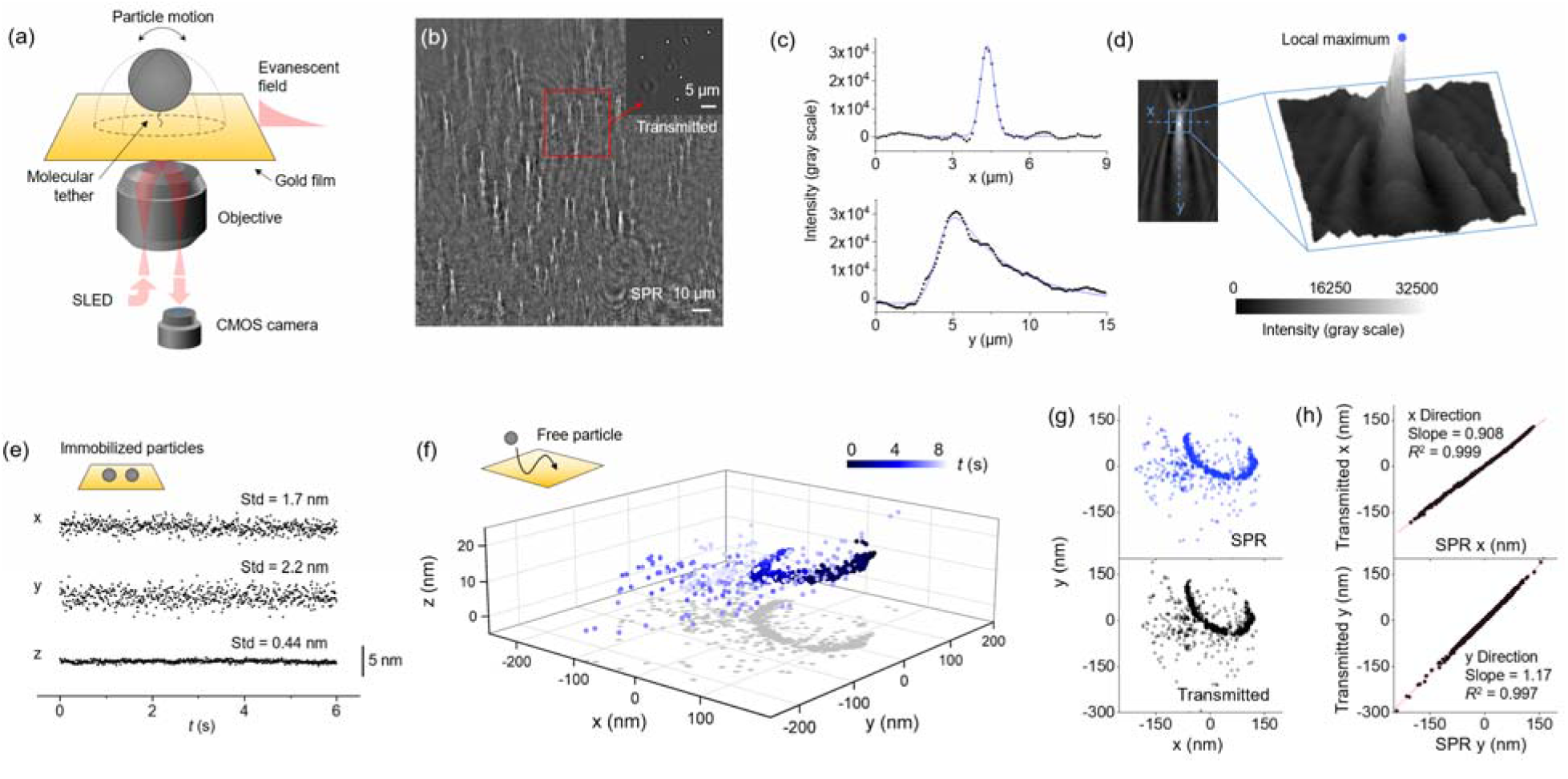
3D tracking of particles using surface plasmon resonance microscopy. (a) Experimental setup. Particles are tethered to a gold surface by single-molecule tethers. Incident light is directed to the surface via a microscope objective to excite SPR. The plasmonic images of the particles are collected by the camera in real-time. (b) A SPR image showing 1 µm PS particles tethered by 16 nm dsDNA. The inset is transmitted image of the squared region. (c) Intensity profiles along x and y directions of the SPR pattern of a 1 µm PS particle shown in d (blue dashed lines). The x profile (top) is fitted with Gaussian distribution and y profile (bottom) is fitted with a right-skewed Gaussian distribution. (d) Localizing the particle by finding the local maxima. The left panel shows an image of a 1 µm PS particle. The center region of the pattern (blue square) is presented in 3D (right), where the z axis is image intensity. The local maximum is found (blue dot) and then used for particle localization by TrackMate. (e) Localization precision of SPR tracking. Precision in xy and z directions are determined to be ∼2 nm and 0.44 nm respectively by tracking the relative position between two immobilized particles. The standard deviation (Std) of relative position fluctuation is defined as localization precision. (f) The motion of a streptavidin coated 1 µm PS particle near the surface revealed by SPR 3D tracking. The gold film surface was passivated with MT(PEG)4 to reduce non-specific absorption. The particle showed a c-shaped pattern caused by interaction with the surface followed by random patterns due to Brownian motion. The shadow on xy plane is projection of the 3D pattern. The particle motion was tracked for 8.1 s at 100 fps. (g) Simultaneous 2D tracking via transmitted light. The xy projection of SPR 3D tracking (top panel) and transmitted light tracking (bottom panel) of the same particle show similar patterns. (h) Evaluation of tracking accuracy by comparing the 2D patterns. The x (top panel) or y (bottom panel) coordinates from SPR tracking and transmitted tracking are plotted in the same graph, and both have a correlation factor *R*^2^ > 0.997. The fitted slope (red line) in x and y are 0.908 and 1.17, respectively.

### 3D tracking of DNA tethered particles

To study the dynamics of nanometer-scaled biomolecules, we tethered particles to the gold surface using short DNA molecules and tracked the particle motion in 3D. The 48 bp (16 nm) double-stranded DNA (dsDNA) was functionalized with a thiol group on one end to couple the gold surface, and a biotin on the other end to capture the 1 µm streptavidin coated PS particle. The density of DNA tether on the surface was adjusted by diluting with spacer molecules to ensure that most particles were tethered by one or a few DNA molecules. In a previous study by Visser et al using bright field 2D tracking,^19^ the motion pattern of short rigid dsDNA-tethered particle was correlated with the number of DNA tethering the particle. They found that single, multiple (likely 2 or 3), and many (> 3) DNA tethered particles displayed characteristic circular, triangular/stripe, and spot patterns. Intuitively, the 2D patterns observed by Visser et al should be the projection of 3D motion onto the image plane. To test our hypothesis, we tracked 121 tethered particles within a 70 µm × 70 µm region at 100 fps for 6 seconds and recorded the motion patterns for each particle in 3D. As expected, the 2D projection showed the same three types of pattern due to single, multiple, and many tethers. In addition, introducing the third dimension reveals more information. For example, particle with one tether has a dome-shaped pattern in space (Figures 2a-c and Supplementary Movie 1), which is due to the free rotation of DNA, while particle with multiple tethers shows a section of the dome (Figures 2d-f), because the motion is restricted by the additional tether. Particle tethered by many DNA molecules is confined within a much smaller region, which is also a section of the dome due to extra restriction by excess tethers (Figures 2g-i). More examples are shown in Figure S3, and a movie showing the motion of particles tethered by different number of DNA is shown in Supplementary Movie 2.

**Figure 2.**
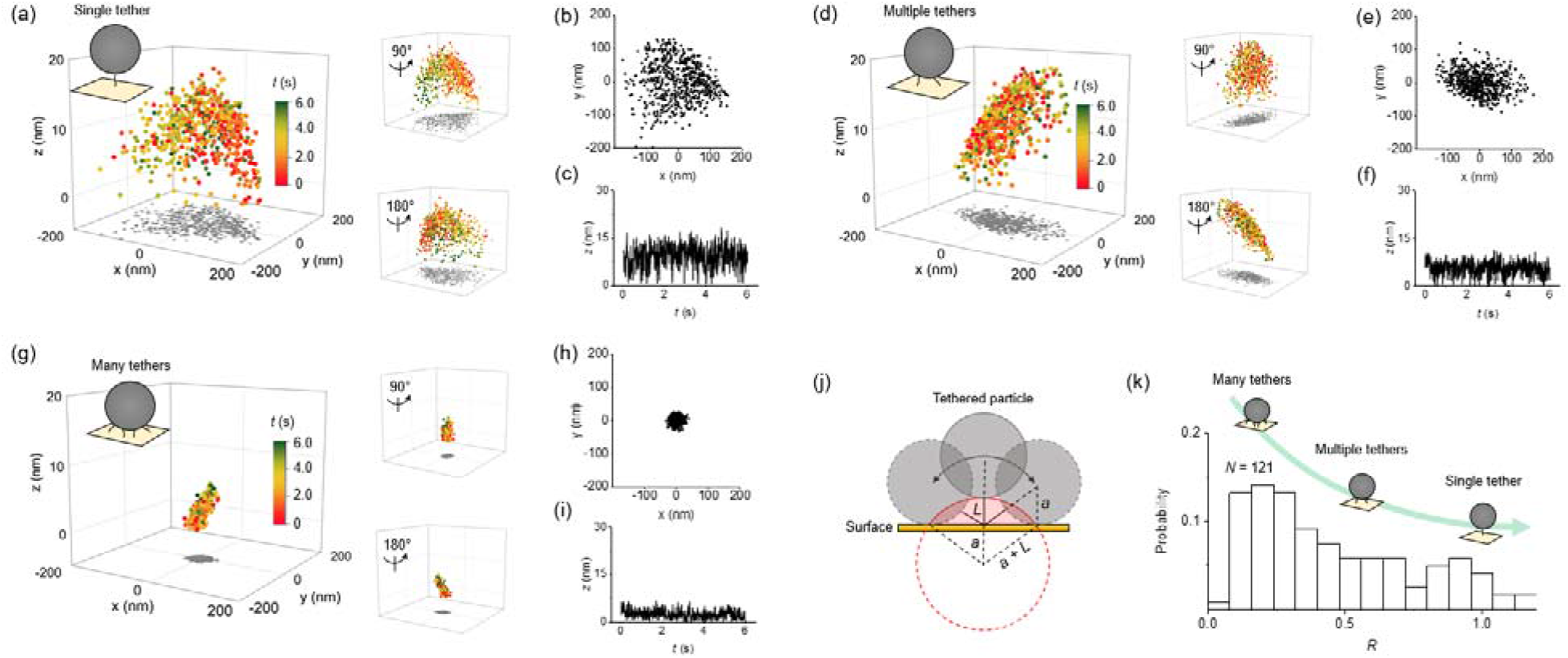
The motion of particle tethered by different number of DNA molecules. (a) The motion of a single DNA tethered particle showing a dome pattern in space. The xy coordinates represent the centroid of the particle, and z coordinate represents the distance from the bottom of the particle to the surface. For clarity, the pattern is rotated 90° and 180° in the right panels. (b) Projection of the 3D pattern onto xy plane. (c) Projection of the 3D pattern onto z-axis. (d) The motion of multiple DNA tethered particle shows a section of dome due to the restriction from additional tethers. The right panels show 90° and 180° rotation of the pattern. (e) Projection of the pattern onto xy plane. (f) Projection of the pattern onto z-axis. (g) The motion of many DNA tethered particle shows that the particle is confined within a small region. The right panels show 90° and 180° rotation of the pattern. (h) and (i) show the projection of the pattern onto xy plane and z-axis, respectively. The tracking frame rate for a, d, and g is 100 fps. (j) Schematic showing a particle with radius of *a* tethered by a DNA with length of *L*. The dome (red solid line and shadow), which is the largest area that the tethered particle can explore, is a section of sphere with radius of *a* + *L*. (k) Distribution of restriction factor *R* obtained from 121 tethered particles. The tether number decreases from many tethers to a single tether as *R* increases from 0 to 1.

The area of particle excursion (*A*) reflects the restriction exerted on particle by tethers. The largest excursion area (*A*_1_) is attained when there is only one tether, which has a spherical cap or dome shape with a radius determined by both particle size and tether length (Figure 2j). Thus, the ratio *R* = *A*/*A*_1_ is a measure of motion restriction with a maximum of ∼1 for a single tether and approaches to 0 for many tethers. We calculated the excursion area for all the 121 particles mentioned above and constructed a histogram in Figure 2k. It can be seen that although the DNA tethering was categorized into 3 states (single, multiple and many), the transition between these states in the histogram is very smooth without obvious peaks, implying that one could not accurately quantify the number of tethers merely from the excursion area. We believe that further combining simulation and experimental data with higher spatial resolution, and using additional parameters such as spatial probability density for comparison may help to solve this problem in the future. Nevertheless, the 3 states criteria can still serve as an acceptable estimation.

### Measuring RecBCD-DNA interaction

The sub-nanometer tracking resolution allows us to probe the intramolecular interaction dynamics of single molecules. To demonstrate this capability, we tracked the interaction between DNA and a DNA enzyme called RecBCD. RecBCD is a hetero-trimeric complex of helicase and nuclease found in *E. coli*, which is responsible for initiating the repair of dsDNA breaks in the homologous recombination pathway.^20^ When RecBCD binds DNA in the presence of ATP, the two helicase subunits unwind the double stranded DNA from one end to the other. Recent single-molecule studies have shown that this unwinding process is accompanied by the rotation of RecBCD due to the double helix structure of DNA,^21^ however, how the rotation is related to the progression of RecBCD on DNA is unclear. This is because by far it is difficult to measure the spatial position and rotation of RecBCD on a single platform simultaneously. DNA length is often measured by magnetic and optical tweezers,^22-24^ while RecBCD rotation is measured by a DNA origami-based method called ORBIT.^21^ Since our tracking technique can probe the 3D coordinates of RecBCD in space, it is possible to obtain DNA length (*L*) and RecBCD rotation angle (*θ*) from the coordinates (Figure 3a).

**Figure 3.**
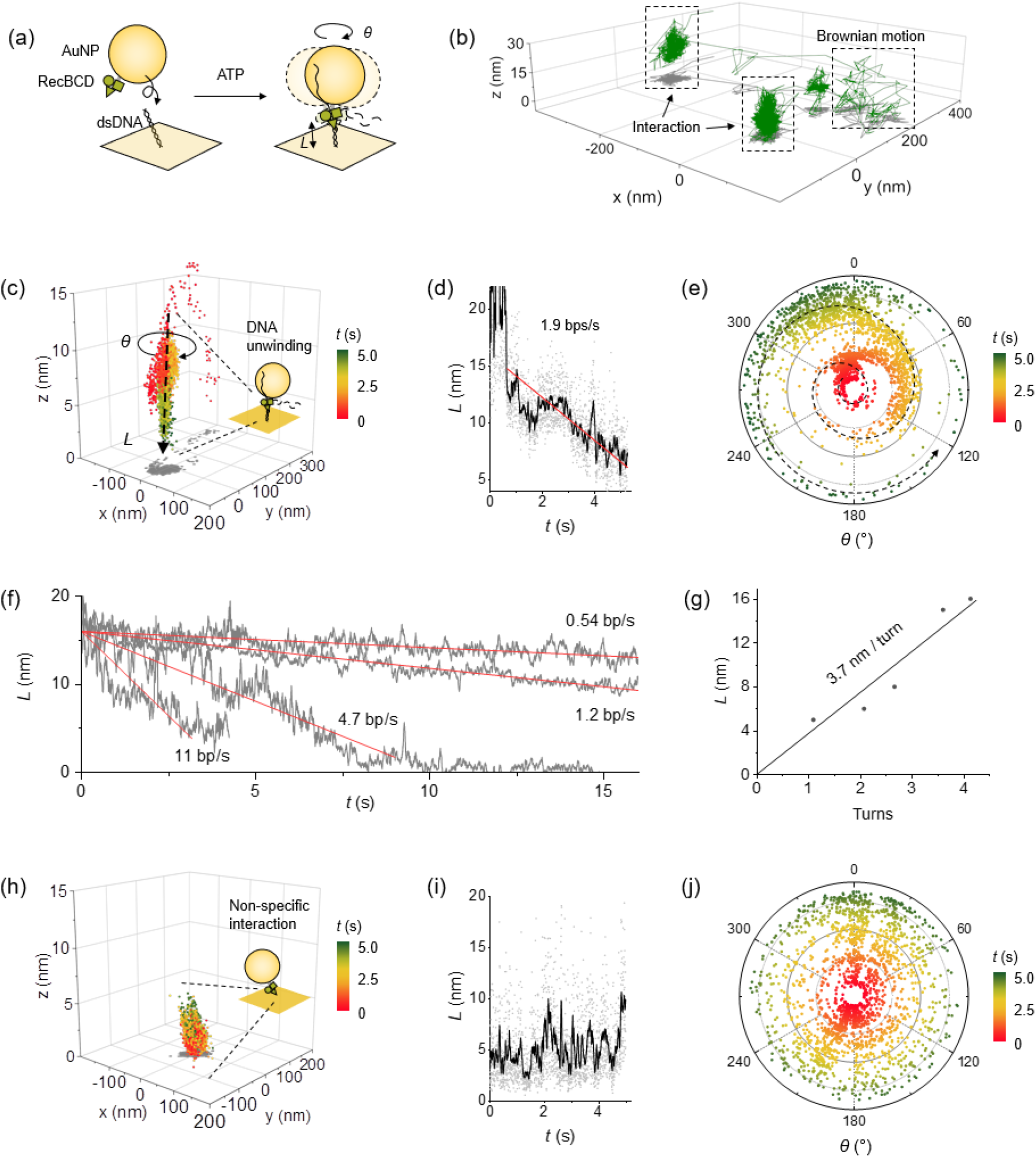
Tracking the interaction between RecBCD and dsDNA. (a) dsDNA is anchored on the gold film, and RecBCD is modified on the surface of a 100 nm gold nanoparticle (AuNP), which unwinds the dsDNA in the presence of ATP. The remaining length of double strand (*L*) and the rotation angle of the RecBCD or AuNP (*θ*) are obtained by tracking the 3D coordinates of the AuNP. (b) Tracking the motion of a RecBCD coated AuNP near the surface. The trajectory of the particle shows Brownian motion and interaction with the surface. (c) Zoom-in of the interaction event shows the motion of AuNP was confined within a nanometer-scaled domain with decreasing *L* and rotating *θ*, which indicates the RecBCD was unwinding the DNA. (d) The change of *L* is obtained from the 3D coordinates in c, where the dots and black curve are raw data and 20 points-smoothed data, respectively. The DNA unwinding rate was determined to be 1.9 bps/s by linear fitting of the curve (red curve). (e) Polar graph showing the rotation of RecBCD upon unwinding. The dashed line marks a possible route of rotation. (f) Unwinding rate of 4 individual DNA molecules. For clarity, the beginnings of the curves are aligned at 16 nm. A total of *N* = 14 DNA molecules were measured, the mean rate was 6.8 bps/s with a standard deviation of 6.5 bps/s. (g) *L* and *θ* (converted to turns) obtained from 5 DNA molecules. The relation between *L* and *θ* were determined by linear fitting of the data, which was 3.7 nm/turn. (h) Control experiment without DNA. The RecBCD coated AuNP bound to the surface via non-specific interaction which showed random fluctuations. (i) *L* change in non-specific interaction, which was calculated using the coordinates in h. The dots and curve are raw data and 20-point smoothed data respectively. (j) Polar graph showing *θ* change in non-specific interaction.

To facilitate the tracking, we immobilized RecBCD on 100 nm gold nanoparticles (AuNPs). Immobilization of RecBCD on surface has negligible effects on enzymatic activity if the RecBCD is properly oriented.^21^ We functionalized the gold surface of the sensor chip with 48 bp dsDNA and filled the sample cell with 1X NEBuffer and 5 µM ATP, which provided a suitable environment for RecBCD to function. This ATP concentration should initiate the unwinding process at a rate of ∼10 bp/s,^25^ which can be readily recorded by the camera at 400 fps. After adding the RecBCD coated AuNPs to the cell, we observed that most AuNPs underwent Brown motion interspersed with transient interactions with the surface, and eventually stuck on the surface. Figure 3b shows the motion of a AuNP within 26.5 s and two interactions events were found. The spatial dimension of each interaction was ∼50 nm in xy and ∼20 nm in z, indicating the AuNP was trapped within a small region. To find RecBCD-DNA interactions, we zoomed in the interaction patterns and plotted the position of the particle against time. For AuNPs with active RecBCD that unwinds the DNA, we found that the AuNP moved unidirectionally from the distal end of DNA to the anchored end on the surface (Figure 3c). The 3D coordinates of the AuNP at different timepoint allowed us to extract the position of RecBCD in contact with the DNA and hence calculate the DNA length change (Figure S4). The length decreased from 14 nm to 6 nm in 4 s after the RecBCD coated AuNP binding to the DNA (Figure 3d). By fitting the length change with a linear model, the reaction rate of RecBCD was determined to be 1.9 nm/s or 6.3 bps/s. Knowing the 3D coordinates also allowed us to extract the rotation of RecBCD around DNA (Figure S4). Figure 3e shows the angular motion of RecBCD during the unwinding process obtained from the coordinates in Figure 3c. The spiral pattern in the polar graph suggests the motion is rotation, and the total rotation angle is 960° (∼2.7 turns) within 4 s. The conversion between DNA length and turns was thus determined to be 3.0 nm/turn or 10 bps/turn, consistent with literature value.^26^ Additional measurements with several different individual DNA molecules (Figure S5) showed that the unwinding rates differed (Figure 3f) but the relation between length and turn was similar (Figure 3g).

To confirm the rotation is due to specific interaction between DNA and RecBCD, we performed a control experiment using a surface without DNA (only with spacer molecules). The RecBCD coated AuNPs diffused to the surface and fluctuated within a small region after hitting the surface (Figure 3h). In contrast to the specific interaction, the non-specific interaction showed a smaller motion range with a constantly fluctuating RecBCD-surface distance of ∼5 nm (Figure 3i), which was likely due to the fluctuation of RecBCD and spacer molecules on the surface. Moreover, the rotation angle of the AuNP was random rather than spiral (Figure 3j). An additional control experiment using DNA coated surface but without ATP showed similar results (Figure S6). All these findings imply that the DNA length change and RecBCD rotation that we observed were due to DNA unwinding. We have studied 135 AuNP-surface interactions from 100 AuNPs, but only 16 unwinding events were found. Such low reaction rate (12%) may arise from the unfavorable orientation of RecBCD immobilized on the AuNPs.

### Identification of specific and non-specific interactions in immunoassay

Besides the DNA, in a more general sense, any molecules or complexes connecting the particle and the surface can act as a tether, and the dynamics of the tether can be probed by tracking the particle. Here we show an example of tracking the particles used in digital ELISA and determine the binding specificity from the motion patterns. Digital ELISA is a recently developed biosensing technique that involves particles as the label for single molecules, e.g. antibodies.^27-29^ We studied the binding of troponin T (TnT), a biomarker for heart diseases,^30^ to its antibody using a sandwich immunoassay provided by a commercial ELISA kit. First, the capture antibody was immobilized on the gold surface via NHS/EDC chemistry. The antibody coverage was carefully controlled to avoid multiple tethers binding to the same particle. Then the surface was blocked by 0.1% bovine serum albumin (BSA) to minimize non-specific interactions. Next, 42 pg/mL TnT was introduced to the system and incubated for 30 min to allow binding to the capture antibody. A second TnT antibody, known as the detection antibody, with a biotin moiety in the Fc domain was used to sandwich the captured TnT. The antibody-antigen-antibody complex was tagged with 1 µm streptavidin coated PS particle via streptavidin-biotin coupling (Figure 4a). We tracked the motion of the particles and the patterns showed that the motion was confined within ±20 nm in xy and 10 nm in z, consistent with the size of the antibody-antigen-antibody complex (about 20 nm) (Figures 4b and S7). Therefore, we infer the motion is due to the specific binding of TnT. The patterns here share some similarities to those of the DNA tethered particles in Figure 2, but the tether numbers cannot be estimated from the patterns, because the model is only valid to rigid tethers. The structure of antibody contains hinges connecting the Fab and Fc domains, which offer flexibility to the structure.^31^

**Figure 4.**
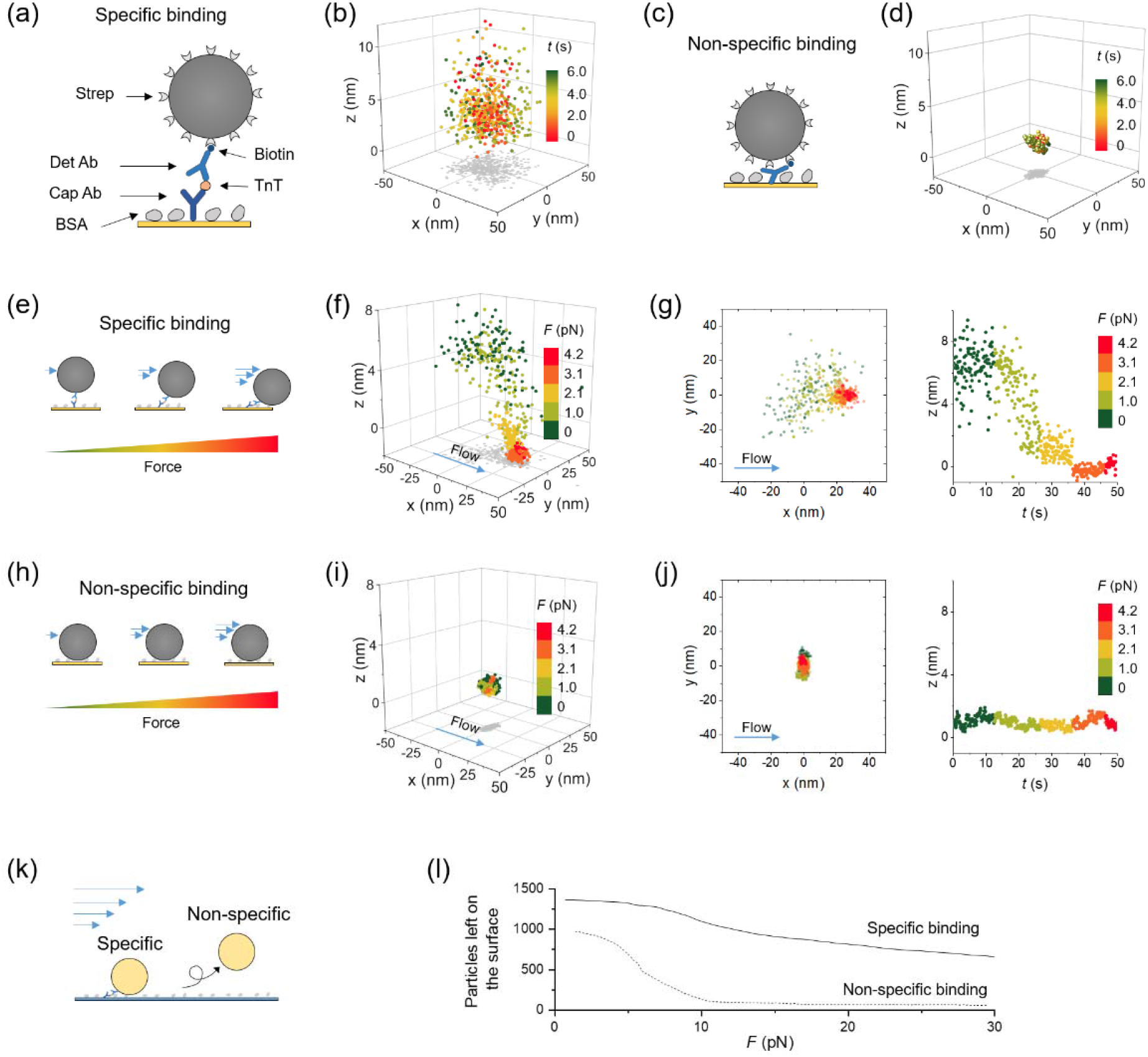
Particle motion reveals the specific binding and non-specific binding of troponin T. (a) Specific binding of troponin T (TnT) in a sandwich immunoassay. TnT is captured by the capture antibody (Cap Ab) immobilized on the surface. The detection antibody (Det Ab) binds to the captured TnT with the Fab domain and captures the 1 µm streptavidin (Strep) coated PS particle via the biotin moiety on the Fc domain. Note that the schematic is not drawn to scale. (b) 3D motion pattern of a representative particle tethered by the antibody-TnT-antibody complex. (c) Non-specific binding in the absence of TnT. (d) 3D pattern of a non-specifically bound particle showing restricted motion. (e) The specific bonding is flexible and can be modulated by a laminar flow. (f) The motion pattern of a specifically bound particle under four different flow rates or forces. (g) Projection of the pattern in f on xy plane and z axis. (h) The non-specific bond is less flexible and cannot be stretched by the flow. (i) Motion pattern of the non-specifically bound particle in flow. (j) Projection of the pattern in i on xy plane and z axis. (k) Further increasing the flow rate ruptures the tether. Tethers with specific interactions are more difficult to break than those with non-specific interactions. (l) Non-specifically bound particles are almost removed at 10 pN, whereas over 50% specifically bound particles remain on the surface at 30 pN. The particles used here were 150 nm gold nanoparticles and the surface was glass. Camera frame rate in b, f, i, and l is 10 fps.

To confirm the particle motion was due to specific binding rather than non-specific adsorption, we performed a control experiment without TnT. The gold surface was functionalized with capture antibody and blocked with BSA, followed by incubation with detection antibody and then the particles. The antibody-antigen-antibody tether could not be formed in the absence of TnT, thus any particles attached to the surface should be attributed to non-specific interactions (Figure 4c). We tracked the motion of these particles and the results are shown in Figures 4d and S7. The patterns are notably smaller than those in the specific interaction, indicating the presence of strong restriction from multiple binding sites, which is a feature of non-specific binding. The motion of specifically and non-specifically bound particles are also shown in Supplementary Movie 3.

To further investigate the difference in tether flexibility, we applied force to the tethered particles using a laminar flow. The flow was generated in a polydimethylsiloxane (PDMS) microchannel mounted on the gold surface. The magnitude of the applied force was controlled by adjusting the flow rate (Figure S8). Four different forces from small to large were applied to the particles. For the specific interaction, the particles were stretched towards the direction of the flow (Figure 4e). Also, as the force increased, the tether became more tightly stretched and the fluctuation diminished (Figures 4f-g and S9). By contrast, the motion of the non-specifically bound particles was not affected much by the force (Figures 4h-j). As we further increased the flow rate, both specific and non-specific bonds were ruptured, but at different flow rates (Figure 4k). We found that over 90% non-specifically bound particles were ruptured at 10 pN, whereas 50% specifically bound particles remained on the surface even at 30 pN (Figure 4l and Supplementary Movie 4). This observation reflects that specific binding is stronger than non-specific binding in terms of binding force, even though the non-specific binding has multiple binding sites. Note that the sealing between gold surface and PDMS is not strong enough to withstand the high flow rate, so we performed the experiment on a glass surface using similar surface chemistry (see Methods). Both the tether flexibility and the rupture force confirm that only specifically bound particles exhibit dynamic motions.

## Discussion

The current setup has a temporal resolution of up to 1000 fps, which is limited by the speed of camera. In a shot noise-limited system, the localization precision is a function of photon number scattered by the particle,^4^ which can be improved by increasing the incident light intensity. At 1000 fps, ∼1 nm precision in xy and ∼0.1 nm precision in z can be readily achieved with the 15 mW SLED light source (Figure S1). We anticipate that using a faster camera and a brighter light source can further improve the spatial resolution to sub-nanometer in all three dimensions at ∼100 µs frame rate, which will enable us to investigate protein conformation change and single base pair change in DNA. However, in a force-free system, the localization precision is limited by molecular thermal fluctuations (Figure 3i), which is a few nanometers depending on the molecular size and flexibility. Stretching the molecule using magnetic or optical tweezers can reduce the fluctuation, but the effect on molecular dynamics should be considered as well.^21^

SPR tracking shows ∼10% linear deviation in the image plane compared to transmitted tracking (Figure 1g), which may arise from different imaging principles between the two approaches. The whole particle is evenly illuminated in transmitted field. In contrast, the illumination in SPR is not uniform due to evanescent field. Therefore, the SPR pattern is dependent on z distance, which leads to inaccurate fitting by the functions. The deviation may also associate with different fitting methods and parameters. We have provided detailed analysis on tracking methods and deviation in Supplementary Note 1 and Figures S10-12. Besides fitting the intensity profile, another way to track SPR pattern is taking advantage of the spatial information,^32^ however, its localization precision and accuracy remain to be explored. After all, the 10% linear deviation should have little effect in determining the molecular dynamics. If high accuracy is required in certain measurements, combining SPR (z direction) and transmitted imaging (xy directions) using the dual-channel configuration (Figure S2) is an alternative solution (Supplementary Movie 1).

## Conclusions

In conclusion, we have demonstrated 3D particle tracking using SPR with sub-nanometer axial precision and milliseconds time resolution. The axial displacement is directly extracted from the scattered light intensity of the particle, requiring no additional optical components. Using the 3D tracking technique, we have studied the dynamics of short DNA and its interaction with enzyme, and quantified enzyme induced DNA unwinding rate as a function of DNA length change. We have also shown that the specific binding and non-specific binding of antibody can be differentiated by analyzing the motion dynamics. We anticipate SPR 3D tracking technique will expand the understanding of single-molecule dynamics and contribute to the development of single-molecule biosensors.

## Methods

### Materials

The functionalized 48 bp DNA was purchased from Integrated DNA Technologies. Methyl-PEG_4_-thiol (MT(PEG)_4_) was purchased from Thermo Fisher Scientific. 1 μm streptavidin coated polystyrene particles and 150 nm streptavidin coated gold nanoparticles were purchased from Bangs Laboratories and Nanopartz respectively. Gold nanoparticle conjugation kit with 100 nm NHS-activated gold nanoparticles was purchased from Cytodiagnostics. RecBCD enzyme was from New England BioLabs. The troponin T detection kit (Elecsys® Troponin T Gen 5 STAT) and troponin T were purchased from Roche. 1X phosphate-buffered saline (PBS) was purchased from Corning. Deionized water with a resistivity of 18.2 MΩ·cm was used in all experiments.

### Experimental setup

The plasmonic imaging system was built on an inverted microscope (Olympus IX-81) with a 60× (NA 1.49) oil immersion objective. The light source was a SLED (SLD260-HP-TOW-PD-670, Superlum) with a wavelength of 670 nm. The plasmonic image of the particles was recorded with a CMOS camera (ORCAFlash 4.0, Hamamatsu) at up to 1000 frames per second. Simultaneous transmitted light imaging was achieved by installing an image splitter (OptoSplit II, Cairn Research) between the microscope and the camera. The light source for the transmitted channel was the stocking halogen of the microscope with a green filter at wavelength of 480-550 nm (IF550, Olympus). The shear flow was generated in a PDMS channel (cross-section: 600 µm × 25 µm) using a syringe pump (Fusion 100, Chemyx).

### Fabrication of DNA tethered particles

The gold surface was cleaned with ethanol and deionized water twice followed by annealing with hydrogen flame to remove contaminates. A PDMS cell was placed on the gold surface for holding solutions. The 16 nm DNA was adapted from reference^33^ with a sequence of 5’ HS-(CH_2_)_6_ -TAG TCG TAA GCT GAT ATG GCT GAT TAG TCG GAA GCA TCG AAC GCT GAT, where the thiol group was used to bind the gold surface. The complementary strand was modified with a biotin at the 5’ end for capturing the streptavidin coated particles. To immobilize the 16 nm DNA on the surface, a mixture containing 1 nM thiolated single strand DNA and 10 µM MT(PEG)_4_ in PBS was introduced to the PDMS well and incubated for 1 hour. Then the gold surface was washed with PBS and incubated in 10 nM complementary DNA for 1 hour to allow hybridization. After hybridization, the surface was washed with PBS again and incubated with streptavidin coated 1 µm PS particles at a concentration of 10^7^ particles/ml for 30 min. Then the surface was slowly washed with PBS to remove untethered particles while not breaking the tethered ones.

### RecBCD-DNA interaction

RecBCD was conjugated to NHS-activated 100 nm AuNPs using a AuNP conjugation kit (Cytodiagnostics). After conjugation, the non-specific sites were blocked with 10% BSA for 10 min. Then the AuNPs were centrifuged at 400 g for 30 min and the supernatant was removed. 100 µL 1X NEBuffer 4 (50 mM potassium acetate, 20 mM Tris-acetate, 10 mM magnesium acetate and 1 mM DTT, pH 7.9) was used to resuspend the AuNPs (particle concentration is ∼7.68×10^10^/mL). The RecBCD coated AuNPs were stored at 4 °C. The 16 nm DNA substrate has the same sequence as mentioned above. The DNA was immobilized on gold surface using the same protocol and density, except that the complementary strand has no biotin group at the 5’ end. After immobilization, the buffer was switched to 1X NEBuffer 4 to match that of RecBCD. ATP used for initiating the reaction was also dissolved in 1X NEBuffer 4.

### Surface preparation for TnT detection

The cleaned gold surface was treated with a mixture of 1 nM O-(2-Carboxyethyl)-O ′ -(2-mercaptoethyl)heptaethylene glycol and 10 µM MT(PEG)4 in PBS overnight. Then the surface was incubated with 50 mM sulfo-N-hydroxysuccinimide (sulfo-NHS) and 200 mM N-ethyl-N ′ -(3-(dimethylamino)-propyl) carbodiimide hydrochloride (EDC) for 15 min to activate the carboxyl groups. 1.7 nM TnT capture antibody solution was immediately applied to the activated surface and incubated for 30 min. The remaining activated sites were quenched by 1 M ethanolamine at pH 8.5. The surface was then incubated with 0.1% BSA for 15 min to block non-specific binding sites. For TnT binding experiment, 42 pg/mL TnT was introduced to the capture antibody functionalized surface and kept for 30 min to allow binding. Then 1.7 nM biotinylated TnT detection antibody was applied to the captured TnT to form a sandwiched structure. Finally, the streptavidin coated PS particles were introduced and kept for 30 min to bind the biotin groups. For the non-specific binding experiment, after functionalizing the surface with capture antibody and blocking with BSA, the detection antibody was directly added to the system in the absence of TnT, and then incubated with the PS particles for 1 h. Immobilization of TnT capture antibody on glass surface was achieved by silanizing the surface with 1% 3-glycidyloxypropyl)trimethoxysilane in isopropanol overnight followed by incubating with 1.7 nM capture antibody solution for 1 h. Then the surface was blocked with 0.1% BSA for 15 min.

## Data Availability

The data that support the findings of this study are available from the corresponding author upon reasonable request.

## Code Availability

The codes that support the findings of this study are available from the corresponding author upon reasonable request.

## Acknowledgements

Financial support from the National Institutes of Health under Award Number R44GM126720 and R33CA235294 is acknowledged.

## Author Contributions

G.M. carried out the experiments and analyzed the data, Z.W. and Y.Y. helped with data analysis, W.J. helped with troponin detection, S.W. helped with instrumentation, S.W. conceived and supervised the project, and G.M. and S.W. wrote the paper.

## Supplementary Information

is available for this paper.

---

**TOC**

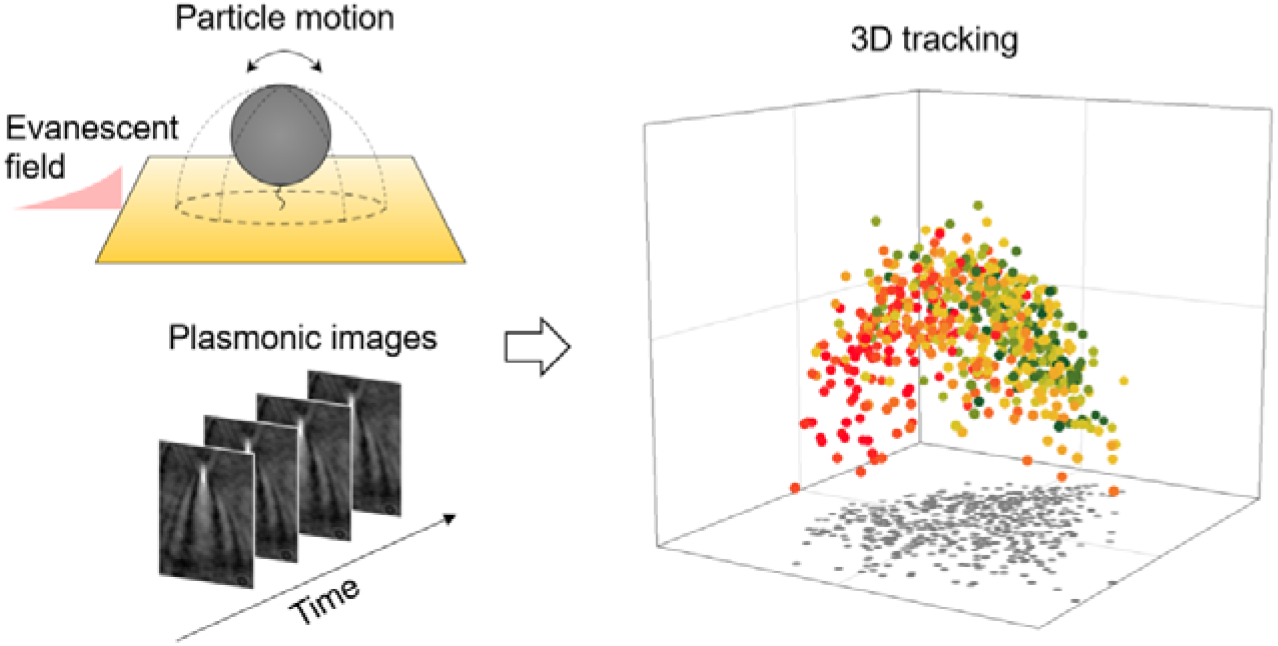

