## Supplementary Information for "Three-dimensional tracking of tethered particles for probing nanometer-scale single-molecule dynamics using plasmonic microscope"

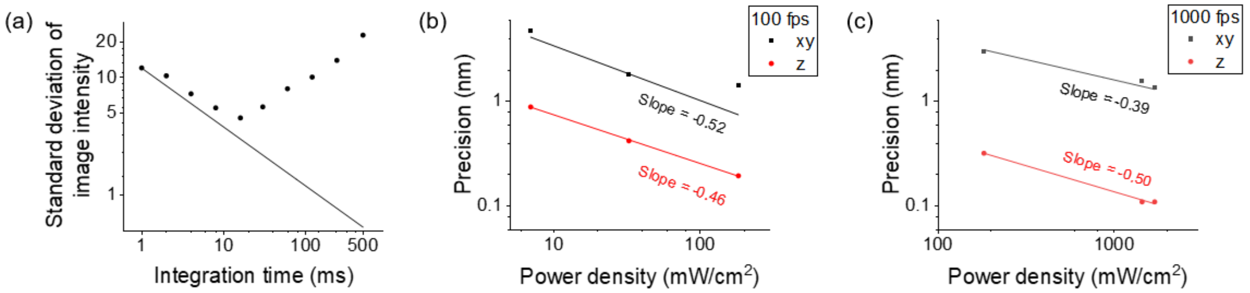


**Figure S1. Localization precision at different incident light intensity and frame rate.** (a) The system was dominated by shot noise at 100 fps or 1000 fps, as determined using a method in literature.[1] Briefly, an image sequence was recorded at 1000 fps for 6 s (incident power density ~1400 mW/cm$^2$) and integrated over different periods to form new image sequences. For each new image sequence, each frame in the sequence was subtracted by its previous frame, and a differential sequence was obtained. The image intensity within a 2.5 µm × 2.5 µm area was measured for each frame in the differential sequence, and the standard deviation of the image intensity was calculated. The standard deviation of image intensity was plotted against different integration time in log-log scale. The solid line is the predicted shot noise level. The plot shows that the system is shot noise limited when integration time is below 10 ms (longer integration leads to drifting). Thus, the tracking we performed at 100 fps and 1000 fps were shot noise dominant. Localization precision in xy (black) and z (red) directions at 100 fps (b) and 1000 fps (c) under different incident power density. Theoretically, the localization precision is inversely proportional to the square root the incident light intensity (or having a slope of -0.5 in the log-log plot) if it is shot noise limited.[2] Linear fits of the data (solid lines) show the slopes for xy and z are close to the predicted value of -0.5, indicating the measurement is shot noise limited.

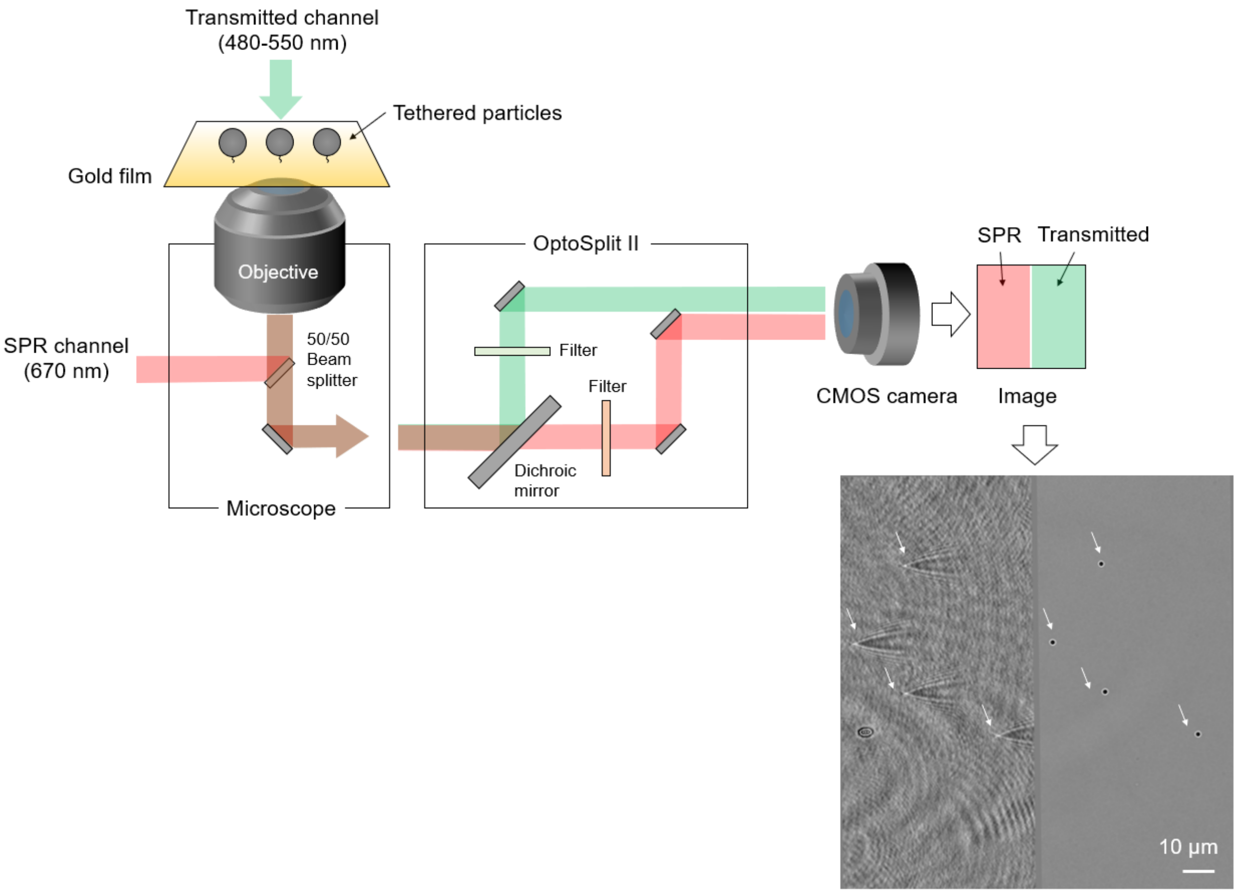


**Figure S2. Simultaneous SPR and transmitted light imaging.** The same region on the gold film was imaged by SPR and transmitted green light in two different channels. A commercial light splitter (OptoSplit II, Cairn Research) was installed between the microscope outlet and a CMOS camera, which separated the two lights by wavelength and aligned them on the imager side-by-side. Particles (size, 1 μm) are marked with arrows.

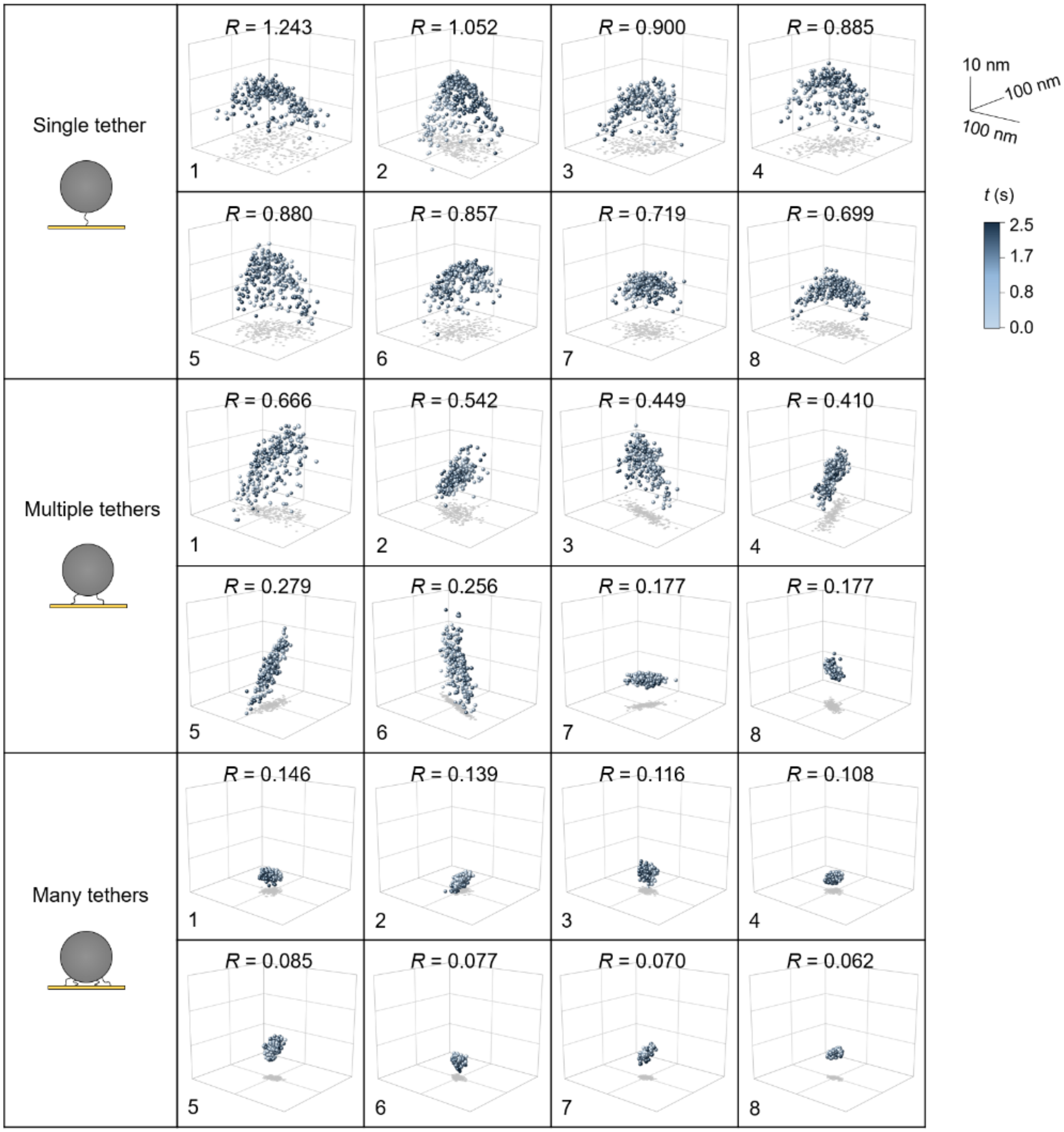


**Figure S3. Additional examples showing the 3D pattern of single, multiple, and many DNA tethered particles.** The *R* value for each pattern is ranked from large to small. DNA length, 16 nm; particle size, 1 µm.

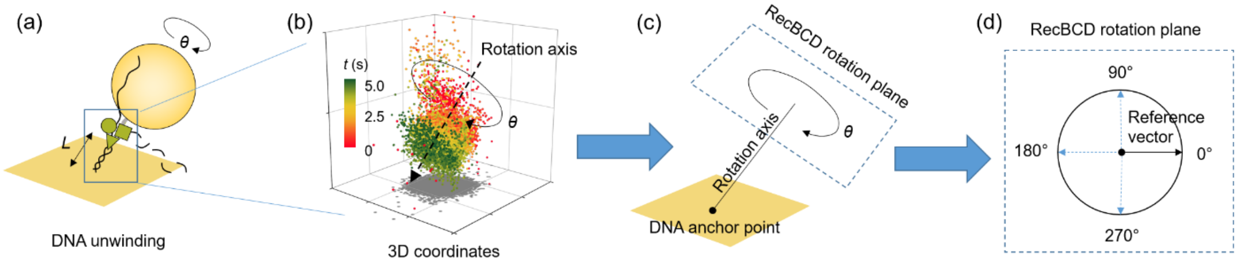


**Figure S4. Extracting DNA length and RecBCD rotation from 3D coordinates.** (a) Schematic showing RecBCD is unwinding a dsDNA. The RecBCD moves from the distal end towards the anchor point on the surface. The remaining dsDNA (or the distance from RecBCD to the anchor point) has a length of *L*. The RecBCD or the AuNP rotates during the unwinding process with an angle of $\vartheta$. (b) 3D coordinates obtained from SPR tracking showing the motion of RecBCD on DNA. (c) The rotation axis is determined by fitting a line through the center of the pattern. The intersection point of rotation axis and $z = 0$ plane is defined as the DNA anchor point. The plane perpendicular to rotation axis is defined as RecBCD rotation plane. (d) The motion pattern is projected to the rotation plane to calculate $\vartheta$ for each point in the pattern. A random vector on the rotation plane with starting point on the rotation axis is set as reference where $\vartheta = 0°$. For each point projected to the plane, a vector connecting rotation axis and the point is created, and the angle between the vector and the reference vector (both on the rotation plane) is calculated as the rotation angle $\vartheta$.

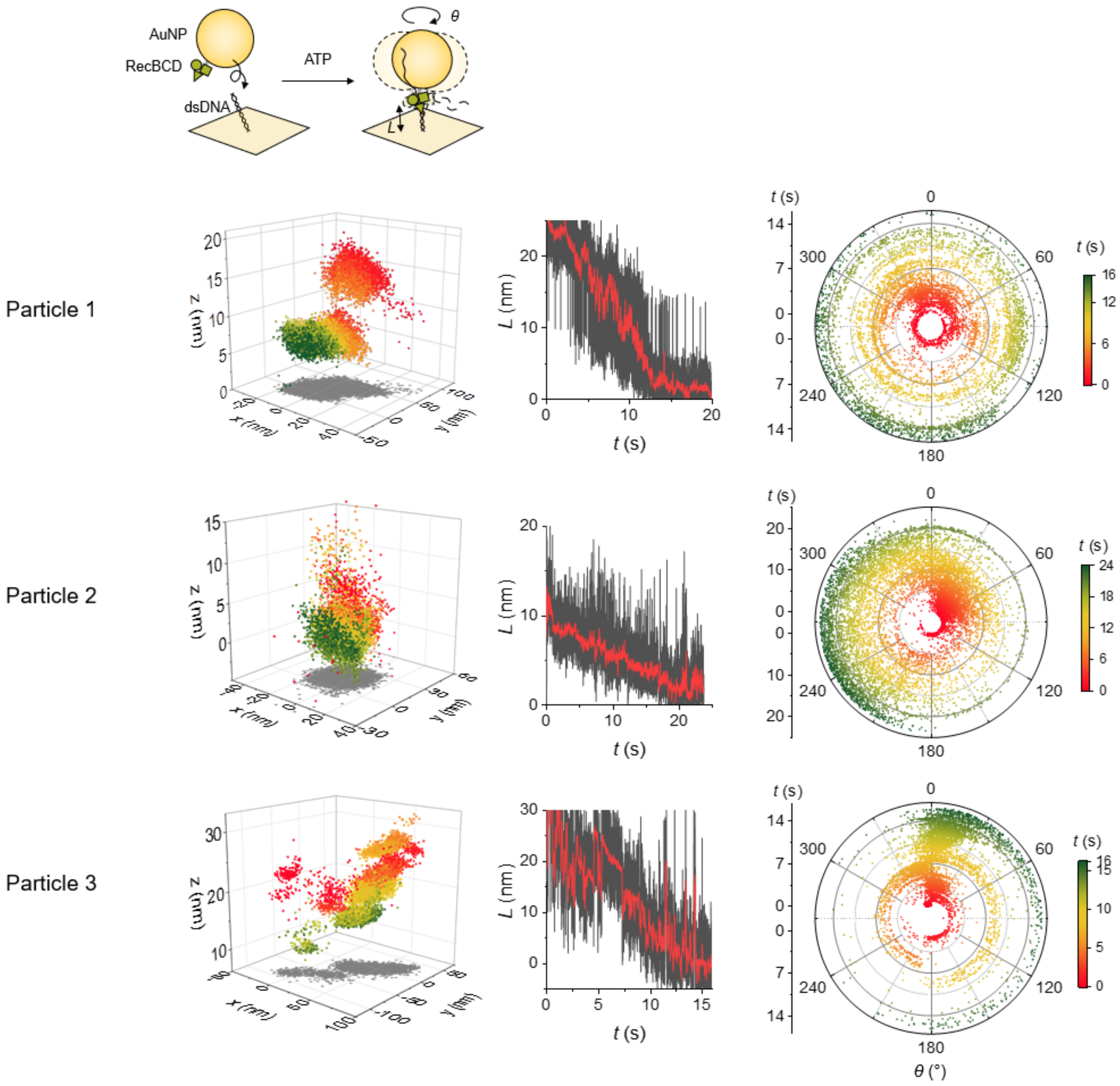


**Figure S5. Three additional examples showing RecBCD unwinds the DNA in the presence of ATP.** From left to right: 3D coordinates of RecBCD, *L* and *ϑ* changes during the unwinding process.

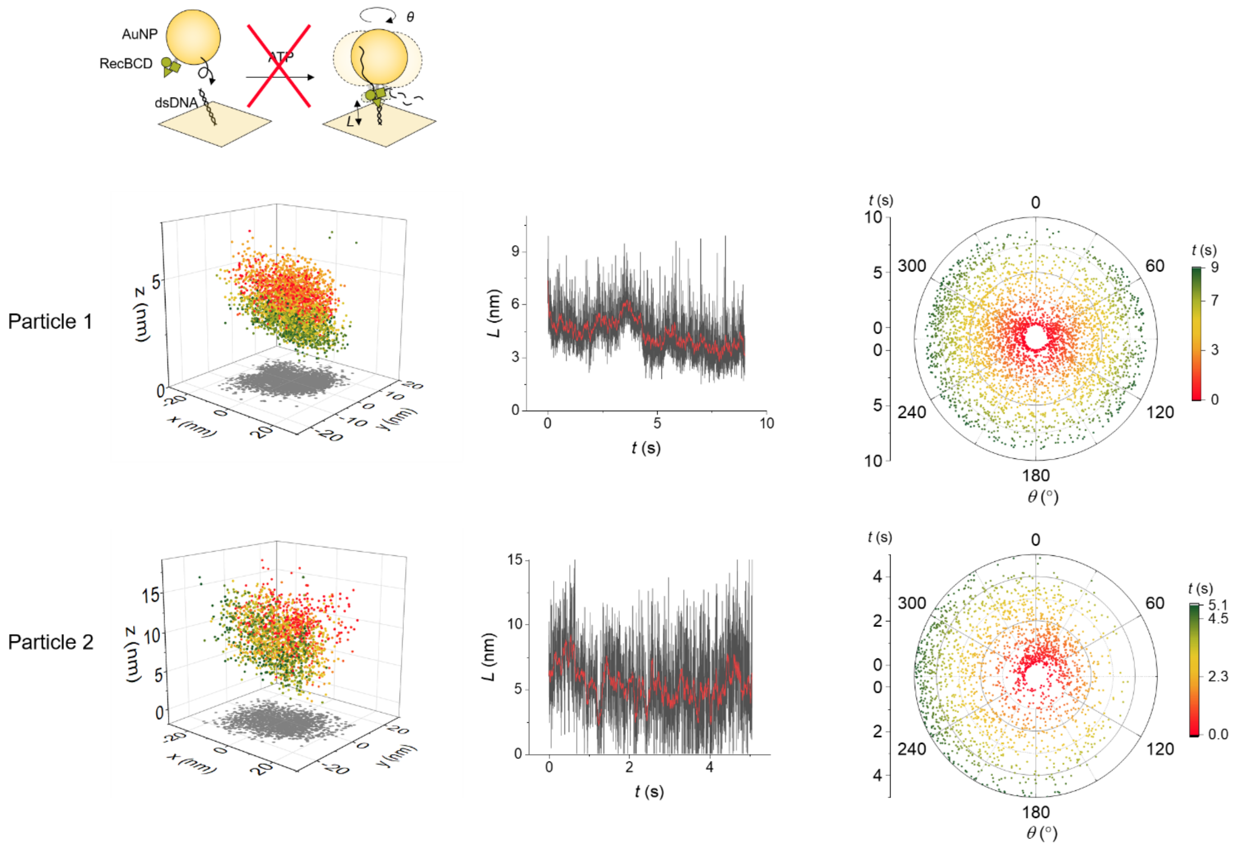


**Figure S6. Control experiment of RecBCD non-specific adsorption without ATP.** From left to right: 3D coordinates of RecBCD, *L* and *ϑ* changes during the unwinding process.

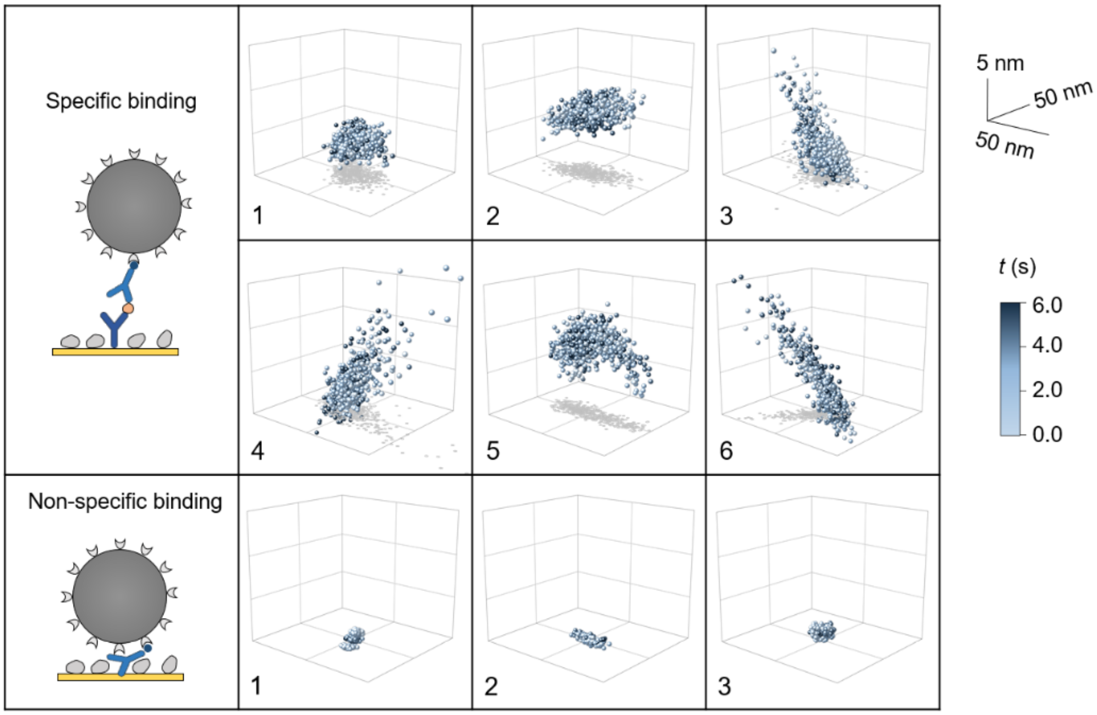


**Figure S7. Additional examples showing 3D patterns of particles tethered by antibody-TnT-antibody complex.** For specific TnT binding (6 examples), the tether consists of a capture antibody, a TnT, and a detection antibody, whereas the non-specific binding has no such tethers and shows more restricted motion patterns (3 examples). The particle is 1 µm streptavidin coated PS particle.

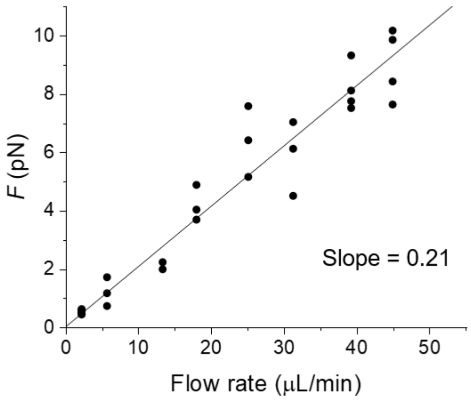


**Figure S8. Calibration of the shear force.** The force is determined by $F = 3\pi\eta Dv$, where $\eta$ is the viscosity of the solution, *D* is the diameter of the particle (150 nm here), and *v* is the linear velocity of the particle. The particle velocity *v* at different flow rates were measured using the camera. PDMS channel has a cross-section of 600 µm × 25 µm. The solid line is a linear fit to the data.

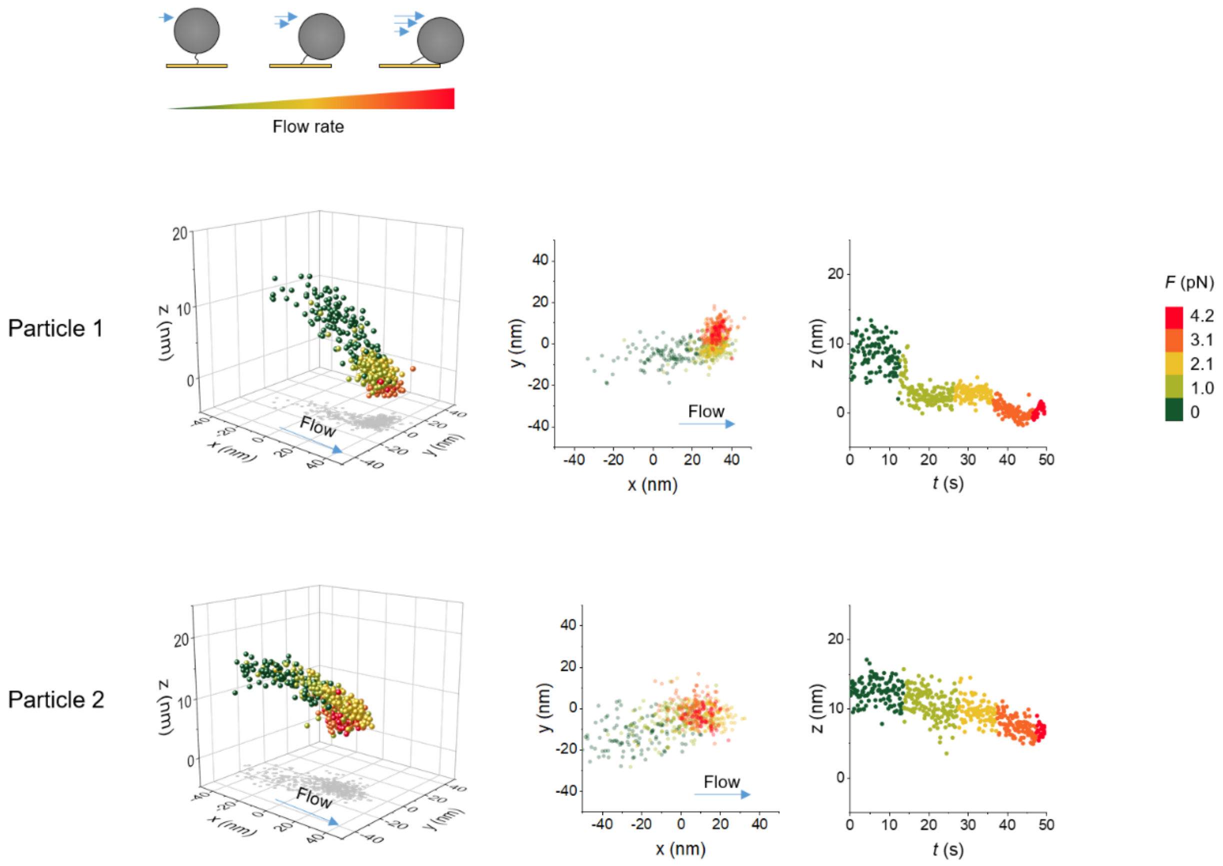


**Figure S9. Two additional examples showing the response of DNA tethered particles in a flow.** Both particles are tethered by multiple DNA molecules, which is determined from the motion patterns. The plots show the motion pattern in 3D, projection on xy plane, and projection on z axis from left to right. Particle size, 1 µm; DNA length, 16 nm.

**Supplementary Note 1**

We compared three different methods for the fitting of SPR pattern. The first one is TrackMate, which uses a LoG (Laplacian of Gaussian) detector to find the local maxima of Gaussian-like spots. The tracking algorithm can be found online.[3] The other two methods are based on 2D Gaussian fitting using MATLAB. The fitted peak position in each frame is used as the particle position. Here we show the detail of the 2D Gaussian fitting. According to Figure 1c, the distribution in x direction is a Gaussian function, given by

$$f(x) = \frac{A_1}{\sqrt{2\pi}w_1} e^{-\frac{(x-x_c)^2}{2w_1{}^2}}. \quad (1)$$

The y direction is a Gaussian skewed by a decay term, which has two components:

$$f_1(y) = \frac{A_2}{t_0} e^{-\frac{y}{t_0}}, \quad (2)$$

$$f_2(y) = \frac{1}{\sqrt{2\pi}w_2} e^{-\frac{(y-y_c)^2}{2w_2{}^2}}. \quad (3)$$

Convolution of the two terms yields the distribution in y direction,

$$f(y) = (f_1 * f_2)(y) = \frac{A_2}{t_0} e^{\frac{1}{2}(\frac{w_2}{t_0})^2 - \frac{y-y_c}{t_0}} \int_{-\infty}^{z} \frac{1}{\sqrt{2\pi}} e^{-\frac{u^2}{2}} du, \quad (4)$$

where the integral is error function, with $z = \frac{y-y_c}{w_2} - \frac{w_2}{t_0}$. The 2D skewed Gaussian function is derived from Eqs. (1) and (4):

$$f(x, y) = f(x)f(y) + A_3. \quad (5)$$

The particle position $(x_c, y_c)$ can be determined by fitting the SPR pattern with Eq. 5. The above equations include a decay term in y direction, which complicates the calculation. To simplify, we can ignore the decay term because the y profile is close to a Gaussian within ~3 µm distance from the centroid peak (Figure 1c), and thus the fitting becomes a 2D Gaussian. We fit the same particle using 2D Gaussian, 2D skewed Gaussian, and LoG detector (TrackMate), respectively, and show the tracking result in Figure S10. The particle motion was tracked simultaneously by SPR and transmitted imaging. By comparing the correlation between SPR and transmitted tracking in x and y direction, we found that the above three fitting methods did not have significant difference, although all of them showed some deviation from the transmitted tracking result.

Next, we explore the origin of the deviation. We found that fitting parameters played a role and could cause distortion of the track. For example, in TrackMate, two initial parameters should be set by user before tracking. Although particle centroid could be successfully localized using different set of parameters, the output tracks were different (Figure S11). We compared the tracks and calculated the correlation with transmitted tracking, and determined the best parameters that gave the largest $R^2$ value. This set of parameters were used for tracking all the particles that captured in the same experiment. For different experiments, it is possible that the best parameters are different, because the SPR pattern is dependent on the focus. For this reason, we recommend that one should determine the parameters for each experiment or fix the focus to avoid the fitting issues.

The tracking deviation caused by fitting parameters is non-linear, because it weakens the linearity of correlation by decreasing $R^2$. Although $R^2$ can approach 1 by tuning the parameters, the slope of correlation is not 1 (Figure S11), which indicates other factors can cause deviation as well. We speculate that the motion in z direction may change the shape of the SPR pattern and hence the fitting result in xy, so we studied the correlation between deviation in xy and z position (Figure S12). We plotted the difference between SPR tracking and transmitted tracking in x and y directions against z position using the data in Figure S10. The slope was close to 0 in x direction but was ~-1 in y direction where the incident light was tilted at SPR angle. The result suggests the y tracking accuracy of SPR image is dependent on z position, likely due to angled SPR illumination light and/or the z dependency of plasmonic wave decay profile.

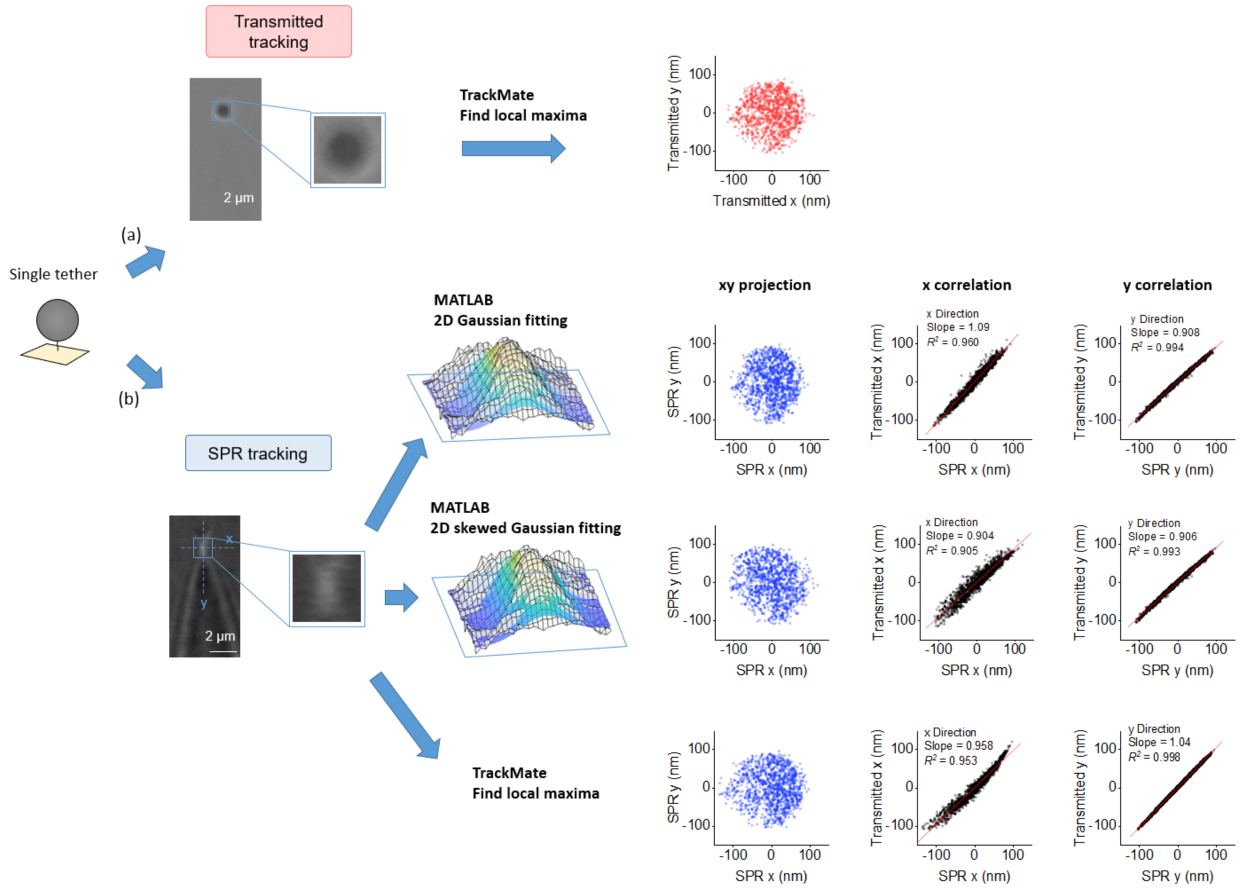


**Figure S10. Different fitting methods for SPR tracking and deviation from transmitted tracking.** The same single DNA tethered 1 µm PS particle was tracked simultaneously by transmitted tracking and SPR tracking. (a) The transmitted images were fitted and tracked by TrackMate and the motion pattern was used to compare with the SPR tracking results. (b) The center region of the SPR pattern was fitted and tracked using three methods. From top to bottom: 2D Gaussian fitting using MATLAB, 2D skewed Gaussian fitting using MATLAB, and finding local maxima using TrackMate. For MATLAB fittings, the image was presented as a 3D mesh, where each grid was a pixel and the height represented image intensity. The colored surface was the fitting result with 2D Gaussian or 2D skewed Gaussian. 3D coordinates were obtained from the SPR fitting methods, but only the xy projection was used to compare with the transmitted tracking result. By analyzing the pattern correlation in x and y directions, we found that the three SPR fitting methods were similar in terms of deviation from transmitted tracking. For ease of operation, we used the well-established TrackMate method in this work.

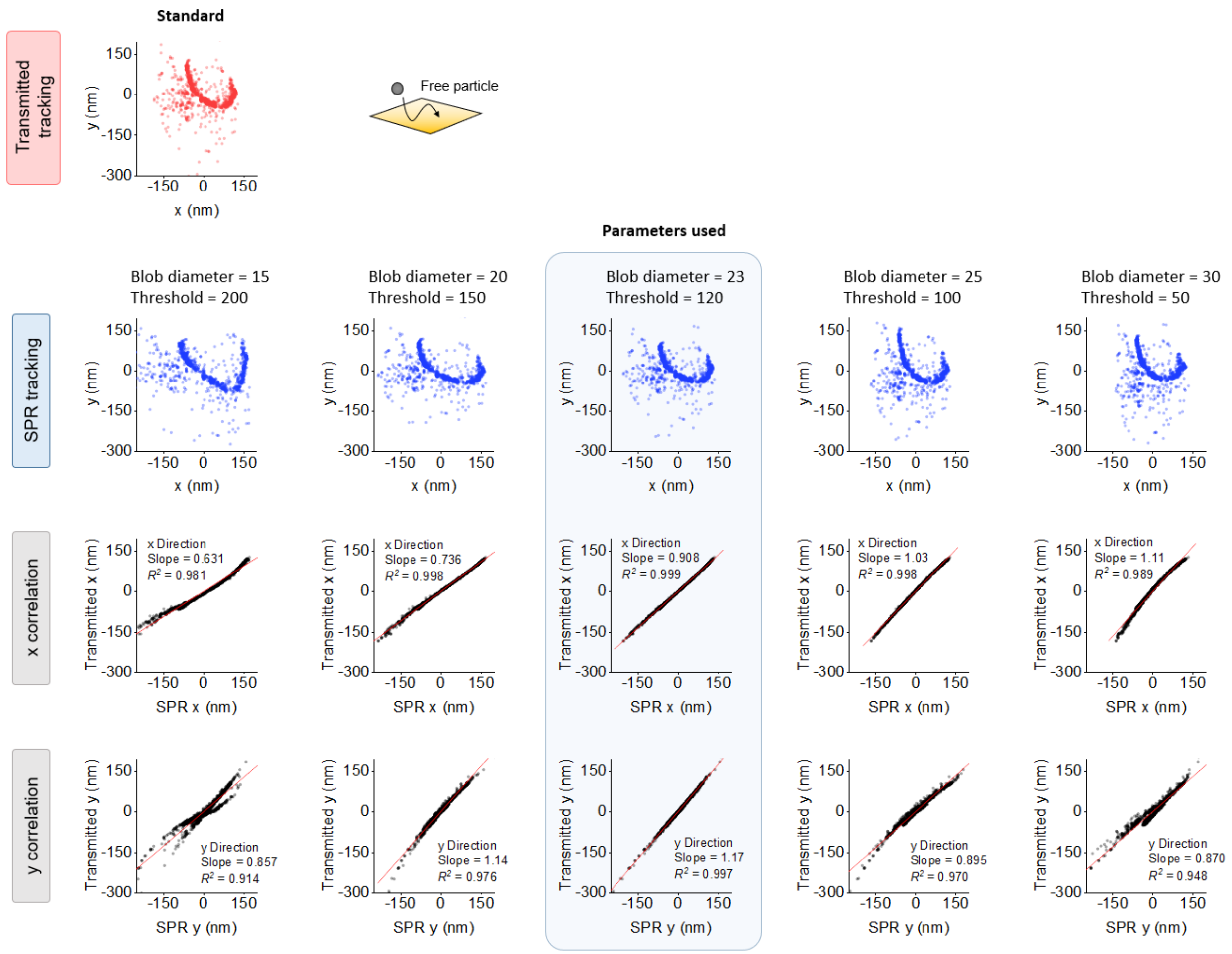


**Figure S11. The effect of fitting parameters on SPR tracking accuracy.** The motion of a free particle (the same one in Figure 1f) was tracked simultaneously using SPR and transmitted light. The transmitted tracking pattern on xy plane was considered as the standard pattern, which was used to study the tracking accuracy of SPR with different fitting parameters. A series of fitting parameters in TrackMate including blob diameter and threshold were tested for SPR tracking (5 sets were shown here), and the correlation in x and y directions were calculated. The parameter set with the highest $R^2$ was chosen for SPR tracking.

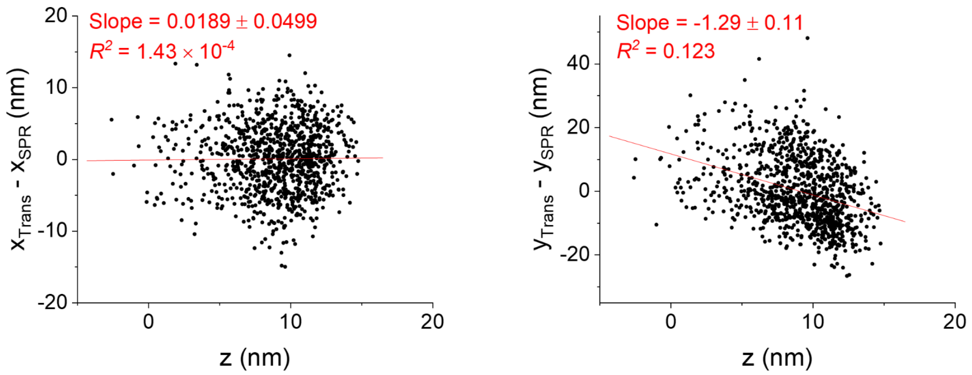


**Figure S12. The tracking deviation in SPR is height dependent.** The difference between transmitted tracking and SPR tracking in x direction (left) and y direction (right) are plotted vs. z, which is obtained from SPR tracking. The slope of the scatter plot in x and y direction are about 0 and -1, respectively, implying that the tracking accuracy is affected by z in y direction (the SPR propagation direction) but not in x direction. The data in the plots are from Figure S10 and tracked by TrackMate.

**Supplementary Movie 1.** Simultaneous SPR 3D tracking and transmitted 2D tracking of the same particle.

**Supplementary Movie 2.** The motion of particles tethered by single, multiple, and many DNA tethers.

**Supplementary Movie 3.** Differentiate specific binding and non-specific binding of troponin T by observing the motions.

**Supplementary Movie 4.** Removing non-specific binding with flow.
